# Strong immunogenicity & protection in mice with PlaCCine: A COVID-19 DNA vaccine formulated with a functional polymer

**DOI:** 10.1101/2023.08.01.551509

**Authors:** Subeena Sood, Majed M. Matar, Jessica Kim, Meredyth Kinsella, Kempaiah Rayavara, Olivia Signer, John Henderson, Joseph Rogers, Bhavna Chawla, Brandon Narvaez, Alex Van Ry, Swagata Kar, Austin Arnold, Jennifer S. Rice, Alanna M. Smith, Daishui Su, Jeff Sparks, Corinne Le Goff, Jean D. Boyer, Khursheed Anwer

## Abstract

DNA- based vaccines have demonstrated the potential as a safe and effective modality. PlaCCine, a DNA-based vaccine approach described subsequently relies on a synthetic DNA delivery system and is independent of virus or device. The synthetic functionalized polymer combined with DNA demonstrated stability over 12 months at 4C and for one month at 25C. Transfection efficiency compared to naked DNA increased by 5-15-fold in murine skeletal muscle. Studies of DNA vaccines expressing spike proteins from variants D614G (pVAC15), Delta (pVAC16), or a D614G + Delta combination (pVAC17) were conducted. Mice immunized intramuscular injection (IM) with pVAC15, pVAC16 or pVAC17 formulated with functionalized polymer and adjuvant resulted in induction of spike-specific humoral and cellular responses. Antibody responses were observed after one immunization. And endpoint IgG titers increased to greater than 1x 10^5^ two weeks after the second injection. Neutralizing antibodies as determined by a pseudovirus competition assay were observed following vaccination with pVAC15, pVAC16 or pVAC17. Spike specific T cell immune responses were also observed following vaccination and flow cytometry analysis demonstrated the cellular immune responses included both CD4 and CD8 spike specific T cells. The immune responses in vaccinated mice were maintained for up to 14 months after vaccination. In an immunization and challenge study of K18 hACE2 transgenic mice pVAC15, pVAC16 and pVAC17 induced immune responses lead to decreased lung viral loads by greater than 90% along with improved clinical score. These findings suggest that PlaCCine DNA vaccines are effective and stable and further development against emerging SARS-CoV-2 variants is warranted.

## Introduction

The COVID-19 pandemic severely impacted public health, the social network, and the worldwide economy. More than 633 million people were infected with SARS-CoV-2, and by the end of 2022 more than 6.6 million people had died globally [1]. While the Director-General of the World Health Organization (WHO) declared in May of 2023 that SARS-CoV-2 is no longer a public health emergency, the WHO’s Strategic Advisory Group of Experts on Immunization has recommended boosters doses. Messenger RNA (mRNA) vaccines such as BNT162b2 from Pfizer/BioNTech and mRNA-1273 from Moderna have been approved with high protective efficacy for COVID-19 [2,3]. These and other approved vaccines targeting SARS-CoV-2, are effective in preventing morbidity and mortality, but have waning effectiveness within a few months [4,5]. In addition, the high rate of SARS-CoV-2 spike protein mutation has led to sequential variants of concern and has resulted in breakthrough infections [6–10]. As the new immune-evading variants of concern emerge, it will be necessary for booster vaccines to be revised quickly.

In addition to developing a SARS-CoV-2 vaccine quickly as a new VOC emerges it will be important the new vaccines are thermostable, easy to manufacture, maintain an excellent safety profile and induce both humoral and cellular immune responses. Importantly, DNA vaccines can induce the desired immune profile, and they also have the advantages of rapid manufacturing in response to new variant outbreaks, a low sensitivity to temperature changes and an excellent safety profile [11–14]. DNA vaccines have been tested in multiple clinical trials [15–18] and demonstrated the induction of immune responses and safety. And a COVID-19 DNA vaccine was approved for emergency use (EUA) in India [19]. However, many of the DNA vaccines being tested are using electroporation [15–18] or needle-free jet devices [20] to increase the efficiency of DNA vaccine delivery. New and easier modes of delivery are being developed.

Alternatives to using devices for DNA vaccine delivery include polymeric nanoparticles, lipid-based delivery systems and viral vectors [21–32]. Nanoparticles have demonstrated increased immunity and expression in a number of studies [28,30]. Liposomes and lipid nanoparticles have also demonstrated the ability to encapsulate DNA vaccines, protecting them from degradation and facilitating cellular uptake [31,33]. Viral vectors for DNA vaccines delivery include viral vectors like adenoviruses or lentiviruses. These vectors efficiently infect cells and deliver the DNA payload, inducing antigen expression and immune responses [34–36]. However, the use of gene delivery via viral vectors has been hampered by problems associated with immune responses targeted to the viral vector itself.

In this manuscript, data is presented from mice immunized with PlaCCine. The DNA vaccine was delivered following formulation with a functionalized polymer and adjuvant. The functionalized polymer was demonstrated to protect the DNA vaccine from degradation. Immunology and protection following vaccination of mice with plasmid DNA vaccines expressing SARS Co-V 2 spike protein for either D614G, Delta or D614G and Delta together is presented. The synthetic delivery system composed of a functionalized polymer was developed and found to enhance the transfection efficiency of formulated plasmid DNA over naked DNA by 5-15-fold in murine skeletal muscle following direct injection. Immunogenicity studies of DNA vaccines expressing spike protein of D614G, Delta, or D614G + Delta combination variants formulated with functionalized polymer and adjuvant resulted in spike-specific humoral and cellular responses as well as protection from viral challenge in K18 hACE2 mice.

## Methods

### Cloning SARS-CoV2 Spike antigens into DNA expression plasmids

To clone the SARS-CoV-2 spike antigens into expression plasmids as illustrated in **Figure 1A**, DNA expressing full length spike with a prefusion-stabilizing 2P modification [37,38] were codon optimized and inserted into the plasmid backbone by Gibson assembly (Transomic Technologies, Huntsville, AL). The spike antigens were from variants D614G (pVAC15), Delta (pVAC17) or both D614G and Delta (pVAC16).

**Figure 1.**
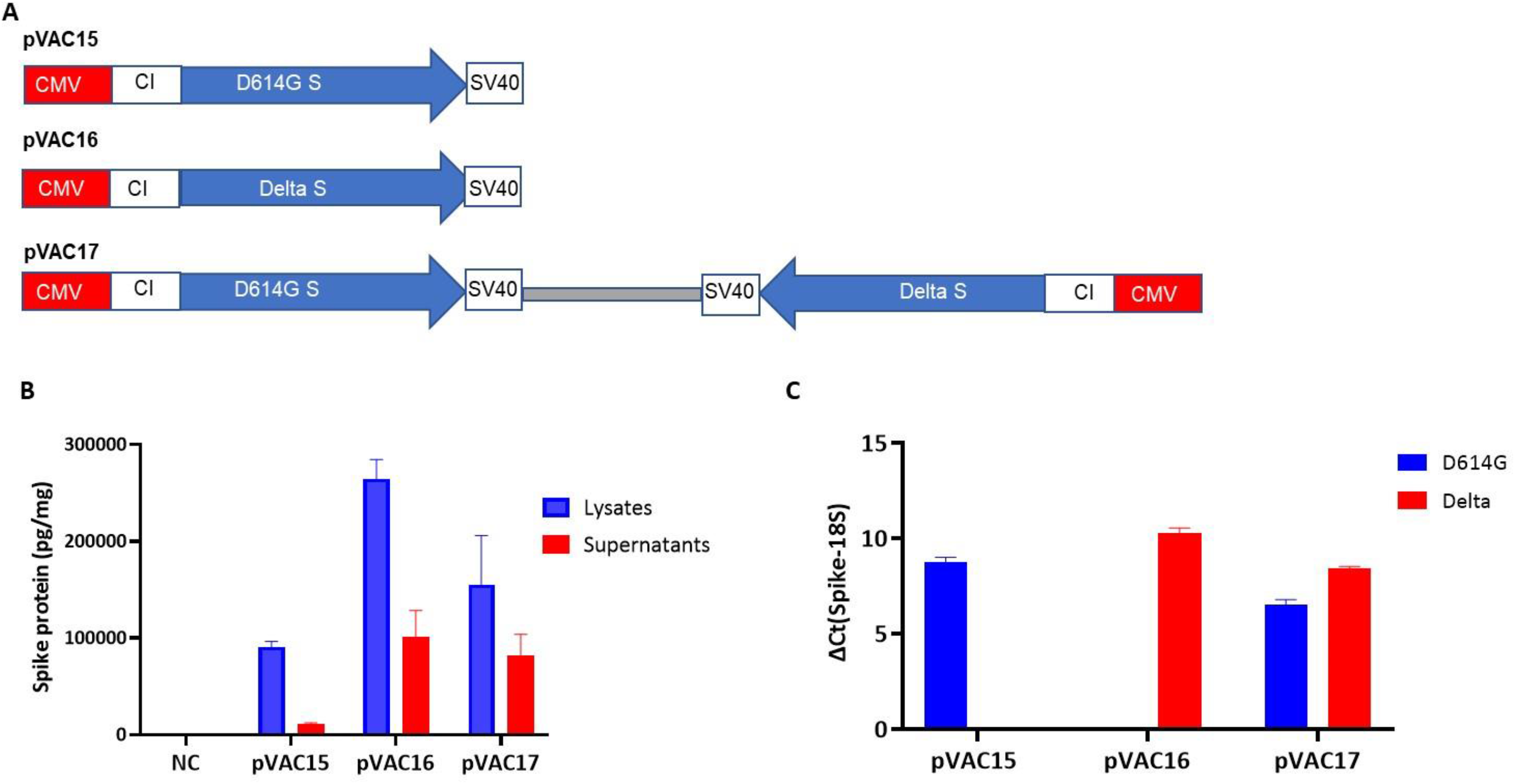
DNA Vaccine construction and expression in mammalian cells. (A) Maps of pVAC15, pVAC16 and pVAC17 are presented. (B) SARS-CoV-2 spike protein expression was determined by ELISA in cell lysates and cell free supernatants of HEK293T cells transfected with pVAC15, pVAC16 and pVAC17. (C) RT-PCR analysis of D614G and Delta spike mRNA produced from prototypes monovalent vaccine plasmids pVAC15 and pVAC16 after transfection in HEK293T cells.

### Luciferase plasmid formulation

Plasmid DNA encoding luciferase gene was diluted in PBS with to a final concentration of 1 mg/mL mixed with adjuvant and functional polymer. The formulation was gently mixed to form a homogenous solution. Ten mice per group were treated via intramuscular (IM) injection bilaterally on day 0. Mice were anesthetized with 2% v/v isoflurane in oxygen. Fifteen minutes before imaging, mice were injected intraperitoneally with 8 mg/mg of luciferin solution (Thermo Fisher). Animals were then anesthetized (2% v/v isoflurane in oxygen) and transferred to the IVIS Lumina LT series III (Caliper Life Sciences). Image acquisition times were kept constant and bioluminescense was measured with a cooled CCD camera.

### Quantification of SARS-CoV-2 Spike protein by sandwich ELISA

To verify the plasmid expressed spike proteins, a sandwich ELISA was developed. In short, 293T cells were grown in DMEM with 10% FBS in 12-well plates and transfected with 200 pM plasmid DNA using OMNIfect transfection reagent for 48 hours. Total protein was isolated by washing cells with PBS and lysing the cells using TENT buffer (10 mM Tris-HCl, 1 mM EDTA, 0.1 M NaCl, 5% [v/v] Triton X100, pH 8.0) for 5 minutes. Lysate was collected and centrifuged at 10,000 rpm at 4°C for 5 minutes. Supernatant was collected and total protein was quantified using Pierce BCA protein assay kit (Cat# 23225, Thermo Fisher Scientific). A 96 well plate was coated with 2 µg/mL spike antibody overnight at 4°C (Cat# 41050-D003, Sino Biological). The plate was then washed with ELISA wash buffer and blocked with 300 µL/well blocking buffer at room temperature for 1 hour and washed again. Standards or sample were then incubated on the plate for 2 hours at room temperature. Following another wash, 100 µL of detection antibody (Sino Biological, Cat# 40150-D001-H) was added at 1 µg/mL for 1 hour at room temperature. After washing the plate, 100 µL/well of TMB substrate was added and incubated in the dark for 20 minutes. The reaction was stopped with 1M phosphoric acid. Optical density was determined using a microplate reader. Expression data is illustrated in **Figure 1B**.

### Measurement of Spike mRNA by Quantitative Real Time PCR (qRT-PCR)

RNA was isolated 24 hours after transfection of HEK293T cells (Qiagen, Cat# 74004). cDNA was synthesized using Verso cDNA synthesis kit (Cat# Ab1453A, Applied Biosystems). qRT-PCR was performed using D614G Spike and Delta Spike gene-specific primers and probes and TaqMan Fast Advanced Master mix (Cat# 4444963, Applied Biosystems) on a QuantStudio 5 system (Applied Biosystem). Expression was normalized to the ribosomal housekeeping gene 18S (**Figure 1C).**

### Mouse immunogenicity studies/animal immunization

Plasmid DNA encoding spike proteins was diluted in PBS and mixed with functionalized polymer and adjuvant. The formulation was gently mixed to form a homogenize solution and used immediately. Female BALB/c mice (6-8 weeks of age) were obtained from Envigo (USA) and housed at Imunon vivarium facility. All mouse experiments were approved by the Institutional Animal Care and Use committee (IACUC) of Imunon. All mice were injected intramuscularly in each of the hind legs with 50 µL on days 0 and 14. Blood was collected on day 13 prior to the second dose. Spleens were removed on day 35 for cellular assays. Blood was collected on day 35 via cardiac puncture and evaluated subsequently by ELISA and neutralization assay.

### IgG antigen S-specific ELISA

Individual serum samples were assayed for the presence of S-specific IgG by ELISA. ELISA plates (96-well, Nunc) were coated overnight at 4°C with 1 µg/mL purified S1+S2 antigen (Sino Biological) in PBS. Plates were washed with PBS and 0.05% Tween-20, followed by blocking with ChonBlock ELISA Buffer (Cat # 90681, Chondrex, Inc.) for 2 hours at room temperature. Plates were washed and serial dilutions of serum in ChonBlock ELISA Buffer were added to the assay plate and incubated for 2 hours at room temperature. Plates were washed and incubated with anti-Mouse IgG (Cytiva) diluted 1:500 in PBS and incubated for 1 hour at room temperature. Plates were washed and TMB peroxidase substrate solution (ThermoFisher) was added to the wells. Reactions were stopped by addition of 1M H_3_PO_4_ (Fisher Scientific) and absorbance was read at 450 nm on an EL808 ULTRA Microplate Reader luminometer (Winooski, VT) with Gen5 ELISA software using a 1 second integration per well. For each serum sample, a plot of optical density (OD) versus the logarithm of the reciprocal serum dilution was generated by nonlinear regression (GraphPad Prism). Endpoint titers were calculated as the reciprocal dilution that emitted an OD exceeding 4 times the background (secondary antibody alone).

### SARS-CoV-2 pseudovirus neutralization assay

HEK293 cells expressing human ACE2 and TMPRSS2a (Invivogen) were seeded at 20,000 cells/well in a gelatin-coated 96-well plate (Corning) for 16 hours at 37°C and 5% CO_2_. In a separate round bottom 96 well plate, mouse sera were serially diluted threefold starting at a 1:10 dilution. A fixed concentration of SARS-CoV-2 GFP pseudotyped virus (BPS Biosciences) was added and the plate incubated for 60 minutes at 37°C and 5% CO_2_. This mixture was then added to the cells and incubated for 48 hours at 37°C and 5% CO_2_. The number of GFP expressing cells in each well was counted on the ELISpot reader (CTL) using the Fluorospot program. Cells infected with only the pseudotyped virus represent 0% neutralization. The IC50 titers were determined using a log (inhibitor) vs. response - variable slope (four parameters) nonlinear function (GraphPad Prism.

### IFN−γ ELISPOT assay

Spleens from immunized mice were collected in sterile tubes containing CTL-test media (CTL, Cleveland, Ohio) supplemented with 1X antibiotic-antimycotic (ThermoFisher Scientific). Cell suspensions were prepared in the MACS Dissociator (Miltenyi). One spleen per C tube was dissociated using the built in MACS protocol. Following dissociation, the cell suspension was strained through a 70 µm MACS smart strainer, placed on a 50 mL tube per C tube. Each C tube including the filter was rinsed with 5 mL of CTL media making the total volume per spleen approximately 10 mL. The cell suspension was centrifuged at 300xg for 10 minutes at room temperature. The supernatant was aspirated, and the spleen cells resuspended in 10 mL of CTL media. The cell suspensions were counted and diluted to 2.5 x 10^6^ cells/mL. Ninety six-well mouse IFN-γ ELISPOT kit (Mabtech, USA) plates pre-coated with anti-mouse IFN-γ capture antibody were rinsed 5 times with PBS to hydrate the membrane and then blocked with supplemented CTL media for at least 30 minutes, prior to addition of cells. Prepared spleen cells were then added to each well at 250,000 cells/well. Each spleen was then stimulated with S1 peptide pool (10-15mers with 11 aa overlap, Miltenyi Cat, #130-127-041) at a final conc of 1 µg/mL for 18-20 hours along with non-stimulated negative controls. PMA solution at the final concentration of 1X was used as a positive control. After stimulation, the plates were washed as per the manufacturer’s instructions. The plates were dried and spots were counted on the Immunospot reader and analyzed with Biospot software (CTL, USA).

### FLOW Cytometry

Flow cytometry analysis was performed to determine if the cellular immune response included both CD8 and CD4 T cells. Briefly, splenocytes were resuspended in CTL test media (Immunospot) and incubated at 37 °C for 2 hours with either no stimulation or S1 peptide simulation (Miltenyi Biotec) at a final concentration of 2 µg/mL before adding GolgiPlug to all samples for 4 additional hours. Following stimulation, cells were stained with Viobility (Miltenyi Biotec) for 15 minutes at room temperature then stained at 4°C for 10 minutes with a surface stain cocktail containing the following antibodies: CD3 PerCP-Vio700, CD4 APC-Vio770, CD8 VB FITC, CD44 PE-Vio770, and CD62L VB423 (all from Miltenyi Biotec). Cells were washed and stained for 10 minutes at room temperature with an intracellular stain cocktail containing the following antibodies: IFN-γ APC and TNF-α PE (all from Miltenyi Biotec). Cells were analyzed on a MACSQuant10 flow cytometer (Miltenyi Biotec). The no-stimulation values were subtracted from the S1-stimulation values for each corresponding sample.

### Mouse Challenge Study

The mouse challenge study was performed at Bioqual. Housing and handling of the animals was performed in accordance with the standards of the AAALAC International, the Animal Welfare Act as amended, and the Public Health Service Policy. Handling of animals occurred in compliance with the Biosafety in Microbiological and Biomedical Laboratories (BMBL), 5th edition (Centers for Disease Control). Pathogen free, 6–8-week-old female B6.Cg-Tg(K18-ACE2)2Primn/J hemizygotes mice were purchased from the Jackson laboratory.

All mice were randomly divided into groups of five. Mice received either placebo, pVAC15 (D614G), pVAC16 (Delta), or pVAC17 (D614G + Delta). Animals were vaccinated intramuscularly with 125 µg/mouse on day 0 and boosted on day 14. The detailed experimental procedure is outlined in **Figure 5**. Groups 1-3 were challenged intranasally (IN) with of SARS-CoV-2 D614G and Groups 4-6 were challenged IN with SARS-CoV-2 Delta variant on day 41. Bioqual utilized the original SARS-CoV-2 USA/NY-PV08449/2020 (D614G) stock, obtained from BEI and expanded at Bioqual (Lot# 091620-230). The stock had a TCID_50_ of 50x10^6^ in hamsters and was dosed neat in mice. The delta stock was obtained from BEI (NR-56116) and expanded at Bioqual (Lot# 70047614). The stock had a titer of 1.4 x 10^6^ TCID_50_/mL and was used at 1:20 dilution. Clinical well-being was measured based on a 0–4 grading system. In the standardized 0–4 grading system, score 0 is normal; score 1 is mild ruffled haircoat and hunched; score 2 is mild disease with ruffled haircoat; score 3 is moderated disease with additional clinical sign such as lethargic, hunched posture, orbital tightening, increased respiratory rate and/or > 15% weight loss; score 4 is showing dyspnea and/or cyanosis, reluctance to move when stimulated, or ≥ 20% weight loss that need immediate euthanasia. All animals were euthanized on study day 48 and collected serum and tissues.

### Lung viral load determination by Tissue culture infectious dose 50 (TCID_50_)

Tissues were weighed and homogenized in 1000 μL of DMEM medium supplemented with 2% heat-inactivated FBS. Tissue homogenates were clarified by centrifugation at 10,000 rpm for 5 minutes and stored at −80°C. Vero-TMPRSS2 cells were plated at 25,000 cells/well in DMEM with 10% FBS were cultured overnight at 37°C, 5% CO_2_ to more than 80% confluency. Medium was aspirated and replaced with 180 μL of DMEM/2% FBS/gentamicin. Twenty microliters of lung homogenate samples were added to the top row in quadruplicate and mixed followed by tenfold dilutions. Positive (virus stock of known infectious titer in the assay) and negative (medium only) control wells were included in each assay setup. The plates were incubated at 37 °C, 5.0% CO_2_ for 4 days. The cell monolayers were visually inspected for cytopathic effect. TCID_50_ values were calculated using the Reed–Muench formula.

### SARS-CoV-2 Neutralization titer by Plaque Reduction Neutralization Assay (PRNT)

D614G (strain details) and Delta (isolate hCoV-19/USA/PHC658/2021) variants were used for PRNT assay. Virus was propagated in Calu3 cells (HTB-55, ATCC) and titrated using TCID_50_ assay on VeroE6 TMPRSS2 cells. PRNT assay was performed using Delta variant. Vero E6 cells (ATCC, CRL-1586) were seeded in 24-well plates at 175,000 cells/well in DMEM/10% FBS/gentamicin. Serial 3-fold serum dilutions were incubated in 96-well plates with 30 PFU of SARS-CoV-2 D614G or Delta strains for 1 hour at 37°C. The serum/virus mixture was transferred to Vero-E6 cells and incubated for 1 hour at 37°C. One milliliter of 0.5% methylcellulose media was then added to each well and plates incubated at 37°C for 3 days. Plates were then washed and cells fixed with methanol. Crystal violet staining was performed and plaques were recorded. IC_50_ titers were calculated as the serum dilution that gave a 50% reduction in viral plaques in comparison to control wells.

### Data Analysis

Statistical analysis of the results and graph creation were done by GraphPad Prism and Microsoft Excel for general statistical calculation, including arithmetic mean and standard deviation. P values of <0.05 were considered significant.

## Results

### Vaccine Formulation and Expression

Luciferase plasmid at 0.1 mg/ml was incubated with non-functionalized and functionalized polymer at concentration of 0.25%, 0.5%, 1.0%, 2.0%, 3.0% and 4.0%. One ug of the DNA mixture was incubated with one international unit of DNase I in digestion buffer for 10 minutes at 37°C. At the end of the incubation period an equal volume of termination buffer was added to the digestion mixture and further incubated for 10 min at room temperature. Formulation of luciferase plasmid (0.1mg/ml) with the functionalized polymer protected the DNA from DNAse degradation as illustrated in Figure 2A at concentrations as low as 0.5% functionalized polymer as evident from gel electrophoresis (**Figure 2A, lane 10)**. The DNA formulated with non-functionalized polymer was completely degraded at all polymer concentrations (**Figure 2A**).

**Figure 2.**
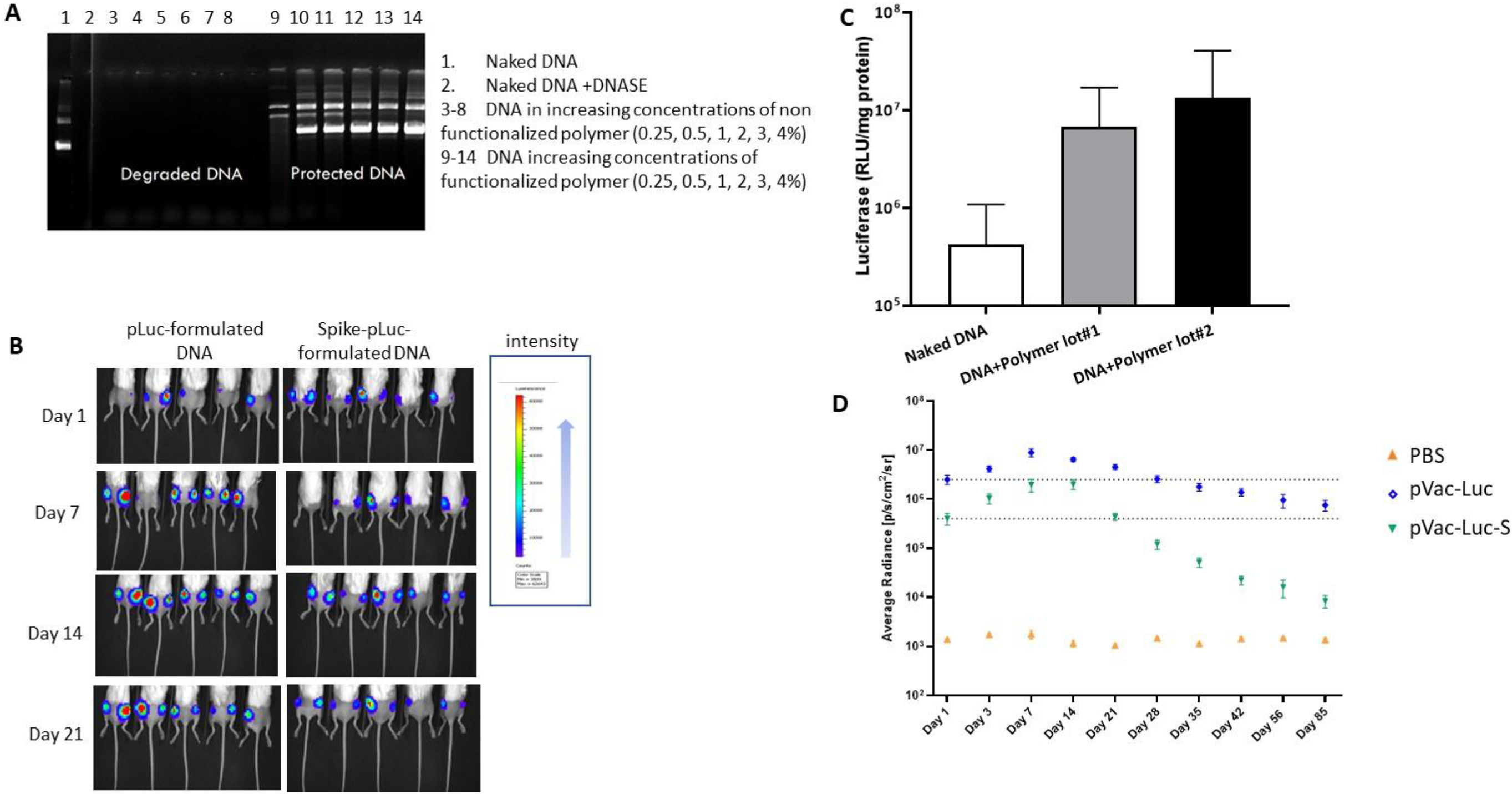
Functionalized polymer protects plasmid DNA from DNase digestion and increases in vivo expression. (A) DNA formulated at a final DNA concentration of 0.1 mg/ml in presence of non-functionalized or functionalized polymer in PBS. One ug of formulated or naked DNA was incubated with one international unit of DNase I in digestion buffer for 10 minutes at 37oC. At the end of the incubation period an equal volume of termination buffer was added to the digestion mixture and further incubated for 10 min at room temperature. Finally, samples were loaded on 1% agarose gel and electrophoresed at 100 V for one hour. (B) Whole mouse luminescence imaging by using In Vivo Imaging System (IVIS) after IM injection with functionalized polymer formulated plasmid DNA encoding firefly luciferase. Luminesce image from individual animals on day 1, 7, 14, and 21 after administration of functionalized polymer formulated plasmid. (C) Expression level of formulated luciferase plasmid with two lots of functionalized polymer and non-formulated (naked) DNA plasmid. (D) Average radiance over 85 days after administration of formulated luciferase or Spike-luciferase plasmid with functionalized polymer and adjuvant.

For gene transfer studies, pLuc was delivered as naked DNA or formulated with functionalized polymer, and luciferase expression was measured by Bioluminescense at days 1, 3, 7, 14, 21, 28, 35, 42, 56 and 85. Data illustrated that DNA formulated with functionalized polymer increased the expression of luciferase by 10 to 15-fold compared to naked DNA (**Figure 2B, C**). A second plasmid expressing luciferase in combination with SARS CoV-2 spike gene was developed as luciferase is not an immunogenic protein. Both vaccines were delivered as described and bioluminescense measured. The bioluminescense decreased more rapidly when the immunogenic spike protein was present. However, bioluminescense was still observed out to day 85 (**Figure 2D**).

### Binding Antibody

To determine if the DNA vaccines formulated with functionalized polymer could induce humoral immune responses, BALB/c mice were immunized by IM administration with either PBS or 125 µg DNA of pVAC15, pVAC16, or pVAC17 on days 0 and 14. Sera samples were collected at days 13 and 35 and tested for binding to spike protein by a standard ELISA. By day 13, all groups demonstrated the formulated vaccine was capable of inducing spike specific antibodies. The IgG end point titers in pVAC15, pVAC16, and pVAC17 vaccinated animals were 625, 5,714, and 764, respectively, on Day 13. Importantly, the PlaCCine formulated vaccine boosted the antibody levels following a second injection. Antibody titers post second injection increased by 3 to 4-fold in all groups. The final mean endpoint titers of pVAC15, pVAC16, or pVAC17 were 232,187, 172,514 and 141,519 respectively (**Figure 3**). While the pVAC16 group had a 10-fold higher antibody titer after the first injection, no difference was observed between the groups in endpoint titers following a second injection.

**Figure. 3.**
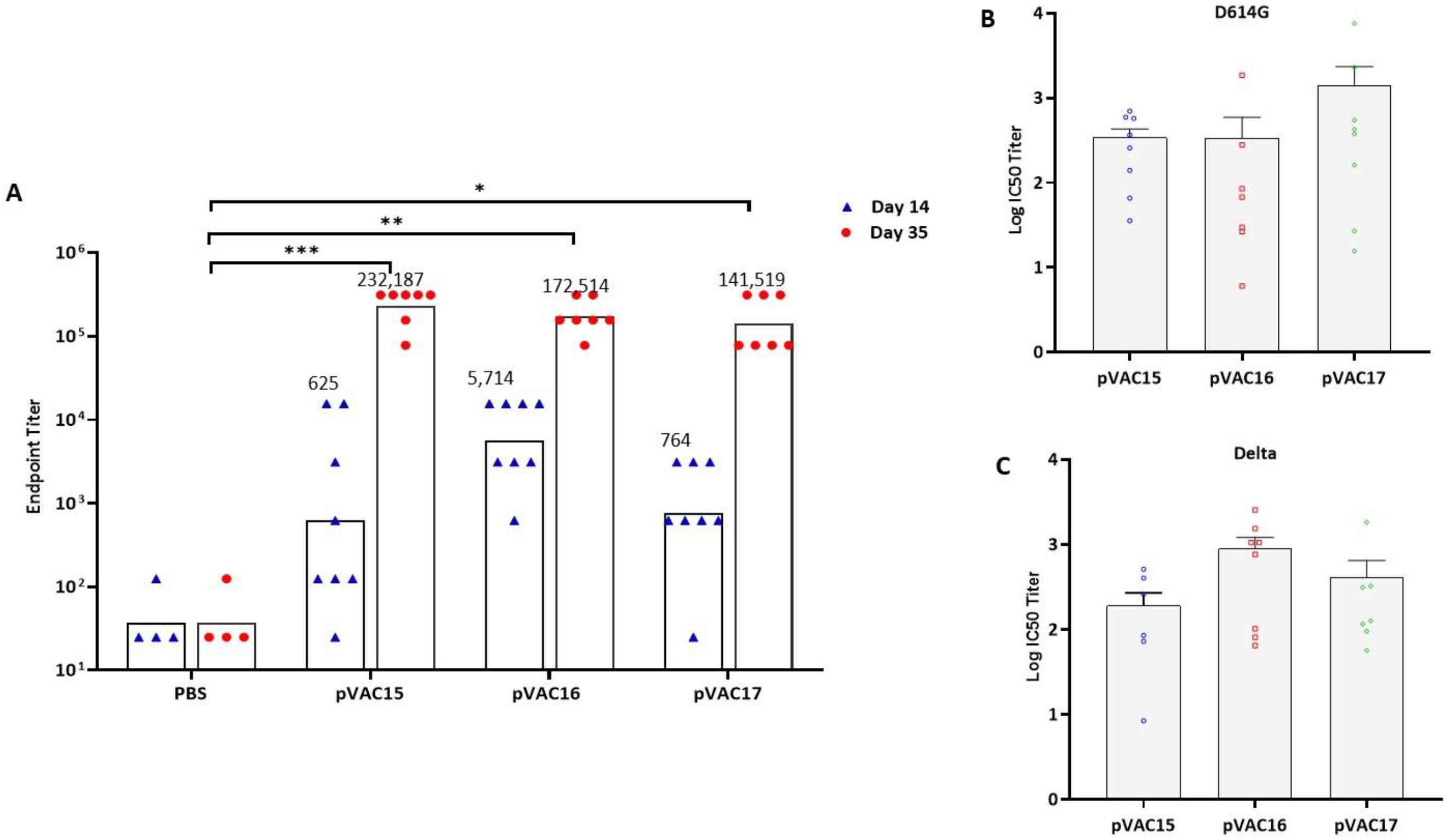
Humoral immune response to pVAC15, 16 and 17 against full length spike protein. (A) Sera from immunized were mice were tested in a standard ELISA. The end point titers to spike full length protein were defined as the reciprocal of the highest dilution that emitted an OD exceeding 4 times the background. (B, C) Neutralization antibodies of pVAC15, 16 and 17 against D614G and Delta. Sera were collected at day 35 from mice immunized on days 0 and 14. Neutralization antibody titers (IC50) were determined for sera from pVAC15, pVAC17 vaccinated mice using D614G variant (B) and Delta (C). GFP pseudovirus and were plotted in Log10 scale. *P< 0.05, **P < 0.01, **** P<0.0001.

### Neutralization Antibody Titer

Induction of neutralizing antibodies is important in the suppression of viral infection and replication. Sera from control and vaccinated BALB/c mice were assessed for neutralization antibody titer with a pseudovirus assay. Evidence of neutralizing antibodies was observed in all vaccinated mice (**Figure 3A, B)**. Neutralization of pseudovirus expressing D614G and Delta was assessed. The mean neutralization against D614G pseudovirus titer (log10 IC_50_, half-maximal inhibitory concentration) was 2.53 for pVAC15, 2.53 for pVAC16, and 3.15 for pVAC17 (**Figure 3A**). The mean log10 IC_50_ nuetralization against Delta pseudovirus was 2.28 for pVAC15, 2.96 for pVAC16, and 2.61 for pVAC17 **(Figure 3B)**. The neutralizing antibody titers from the bivalent pVAC17 (D614G/Delta) vaccine immunized animals were equally effective neutralizing D614G or Delta variants of SARS-CoV-2 while the single variant pVAC15 (D614G) vaccine was less effective against the Delta variant. The single variant pVAC16 (Delta) vaccine was slightly less effective against the D614G variant.

### Cellular and Durability of the Immune Response

The number of splenocytes capable of secreting IFN-γ following *in vitro* stimulation with spike peptides was measured by a standard IFN-γ ELISpot kit. All vaccines demonstrated induction of cellular immune responses following two injections. Evidence of the induction of a combined CD4 and CD8 cellular responses following vaccination with pVAC15, pVAC16, and pVAC17 is illustrated in **Figure 4A**. Five hundred fifty five spot forming cells (SFC) per 10^6^ splenocytes for pVAC15, 660 SFC per 10^6^ splenocytes for pVAC16, and 382 SFC per 10^6^ splenocytes for pVAC17 were observed.

**Figure 4.**
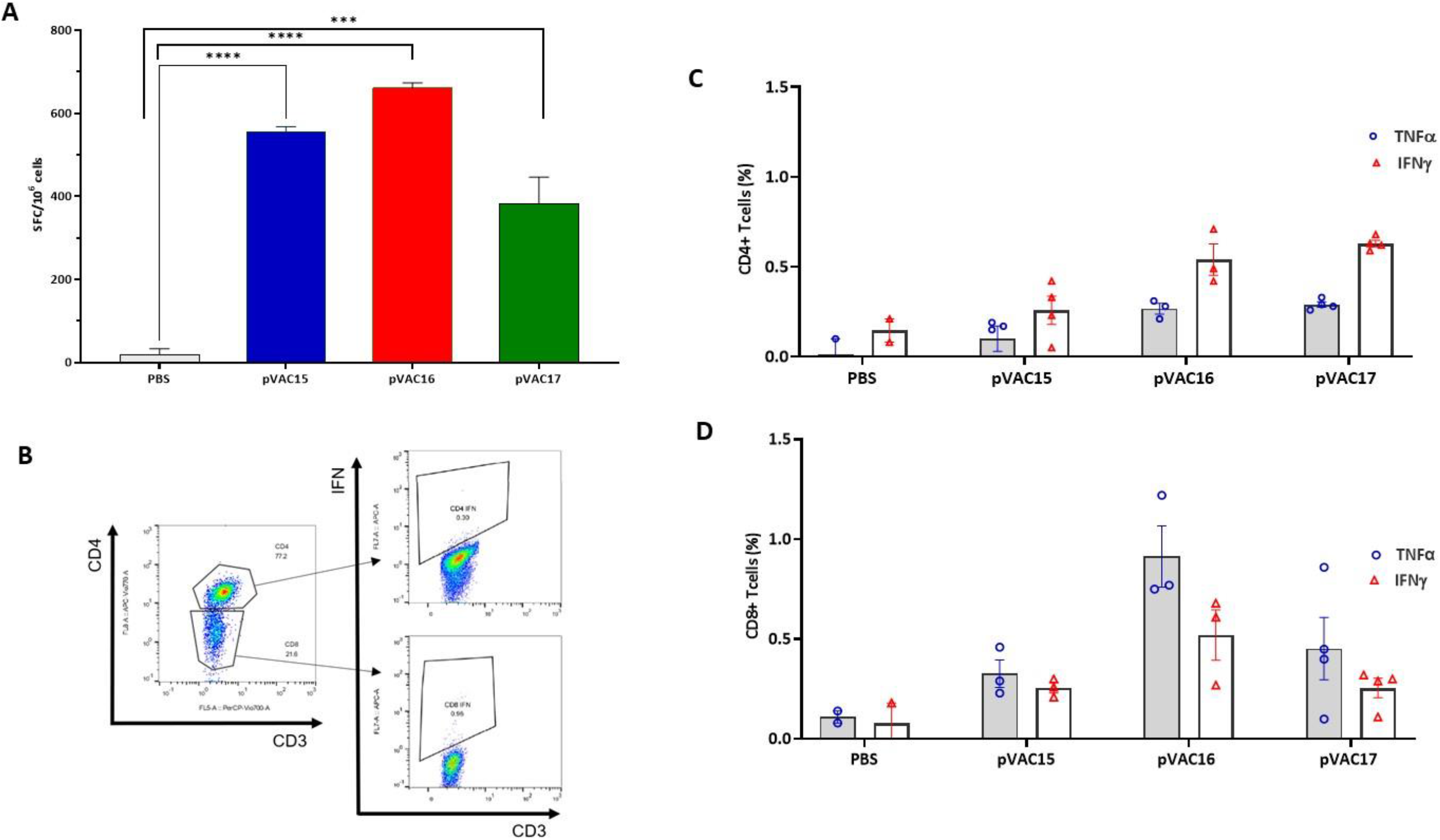
Cellular immune response to pVAC15, 16 and 17. (A) Detection of IFN-γ responses in Balb/c mice post administration of DNA vaccine pVAC15, 16 and 17. IFN-γ responses were analyzed in the animals on day 35. T cell responses were measured by IFN-γ ELISpot in spleenocytes stimulated for 20 hours with overlapping peptide pools spanning SARS-CoV-2 spike, S1 region. The phenotype of the cellular response was measured by flow cytometry. (C) CD4 and (D) CD8 T cells expressing IFN-γ and TNF-a following in vitro spike stimulation are presented. *P< 0.05, **P < 0.01, **** P<0.0001.

In an effort to both distinguish the cellular response as either CD4+ or CD8+ and study the durability of the immune response a second mouse study was designed. BALB/c mice were immunized by IM administration with either PBS or 125 µg DNA of pVAC15, pVAC16, or pVAC17 on days 0 and 14. Sera samples were collected at 1, 2, 3, 5, 6, 9, 12 and 16 months. The spleens from control and immunized animals were collected at **termination of the study (16 months)**. Splenocytes harvested from vaccinated and placebo control mice were stimulated with spike-specific peptides for 6 hours. To determine if both CD4 and CD8 vaccine specific immune response were induced by the PLaCCine formulation, following stimulation cells were stained for CD4, CD8 and stained intracellularly for IFN-γ, and TNF-α. The data illustrated that PlaCCine induced both spike specific production of IFN-γ, and TNF-α in both the CD8+ and CD4+ populations (**Figure 4C, D**), demonstrating a long term cellular memory response.

Similar to the cellular immune response the antibody response remained robust after 12 months post vaccination. However, importantly, while the binding antibody response declined starting at 3 months (**Figure 5A**) the neutralizing antibody response remained constant, **Figures 5 B and C**.

### Protection efficacy in K18-hACE2 transgenic mice

K18-hACE2 mice (n=5) received two doses of 125 µg of either pVAC15, pVAC16, pVAC17 or placebo at two week intervals via IM injection (**Figure 6A**). Four weeks post-immunization, mice were challenged intranasally with either D614G or Delta variant strain (**Figure 6A**). Body weight changes were measured daily for seven days post-infection (dpi). D614G variant virus infection in placebo group showed rapid weight loss with maximum weight loss of 13% whereas pVAC15 vaccinated and pVAC17 vaccinated groups lost 9% and 5% body weight, respectively (**Figure 6B**). Disease clinical score peaked at 6 dpi in the placebo group with mean score of 3. Vaccinated groups showed a clinical score of 3 and 1 at 7 dpi in pVAC15 and pVAC17 groups, respectively (**Figure 6C**). Following infection with D614G virus placebo group had a 20% mortality at 6 dpi. The pVAC15 had a 40% mortality at 7dpi, whereas pVAC17 dual antigen immunized mice had 100% survival (**Figure 6D**). Lung viral load at 7 dpi of 2.6×10^6^ TCID_50_/gm were detected in Placebo group whereas pVAC15 immunized group had more than a 1.5 log lower viral load with 1.0×10^4^ TCID_50_/gm. Two out of 5 mice lungs had no detectable virus; in pVAC17 vaccinated group showed no detectable virus was observed in 4 out of 5 mice with mean virus titer of 3.04×10^4^ TCID_50_/gm (**Figure 6E**).

**Figure. 5.**
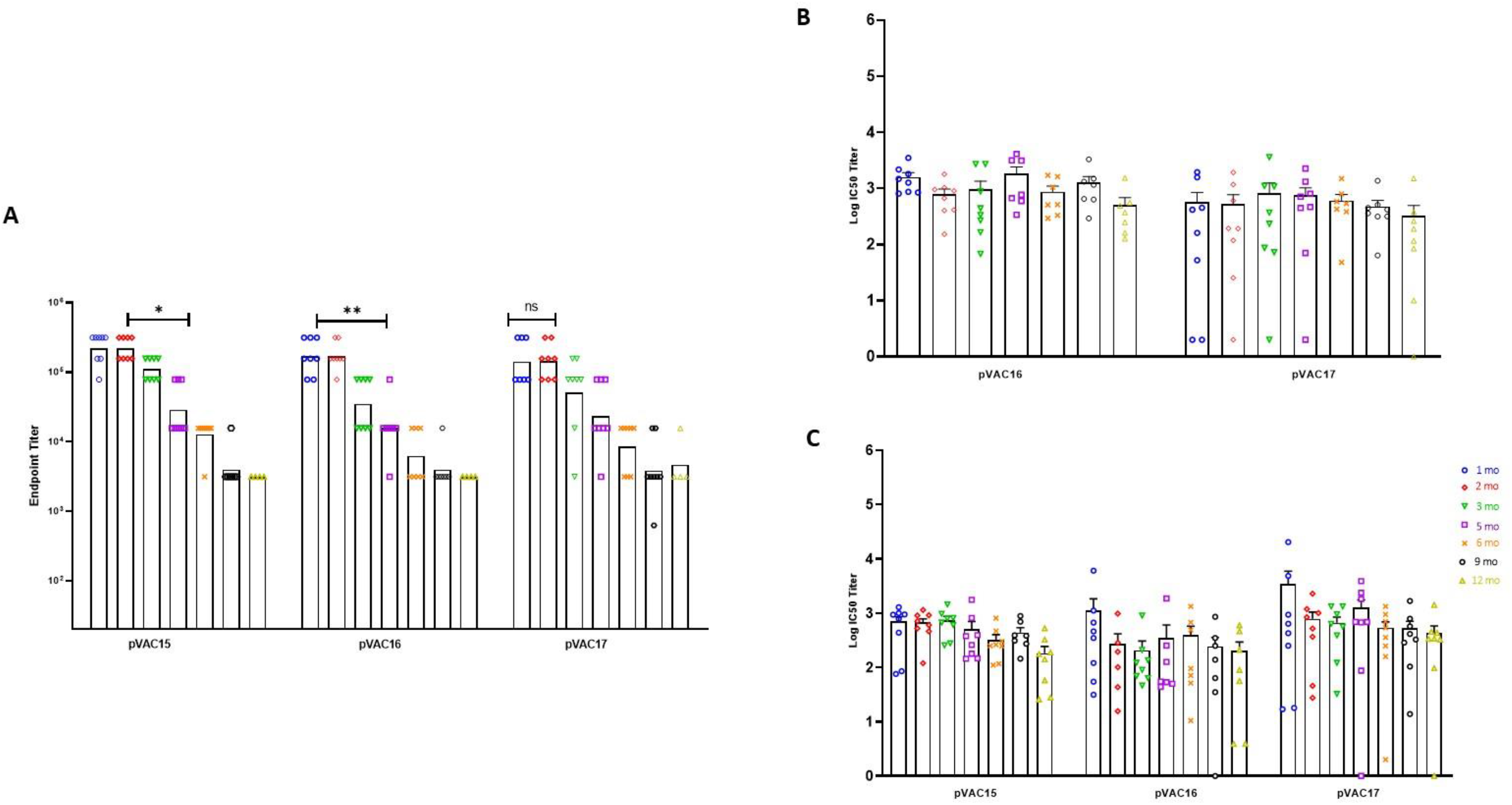
Durability of Humoral immune response to pVAC15, 16 and 17 against full length spike protein. (A) Sera from immunized were mice were tested in a standard ELISA. The end point titers to spike full length protein were defined as the reciprocal of the highest dilution that emitted an OD exceeding 4 times the background. (B, C) Neutralization antibodies of pVAC15, 16 and 17 against D614G and Delta. Sera were collected at day 35 from mice immunized on days 0 and 14. Neutralization antibody titers (IC50) were determined for sera from pVAC15, pVAC17 vaccinated mice using D614G variant (B) and Delta (C). GFP pseudovirus and were plotted in Log10 scale. *P< 0.05, **P < 0.01, **** P<0.0001

**Figure 6.**
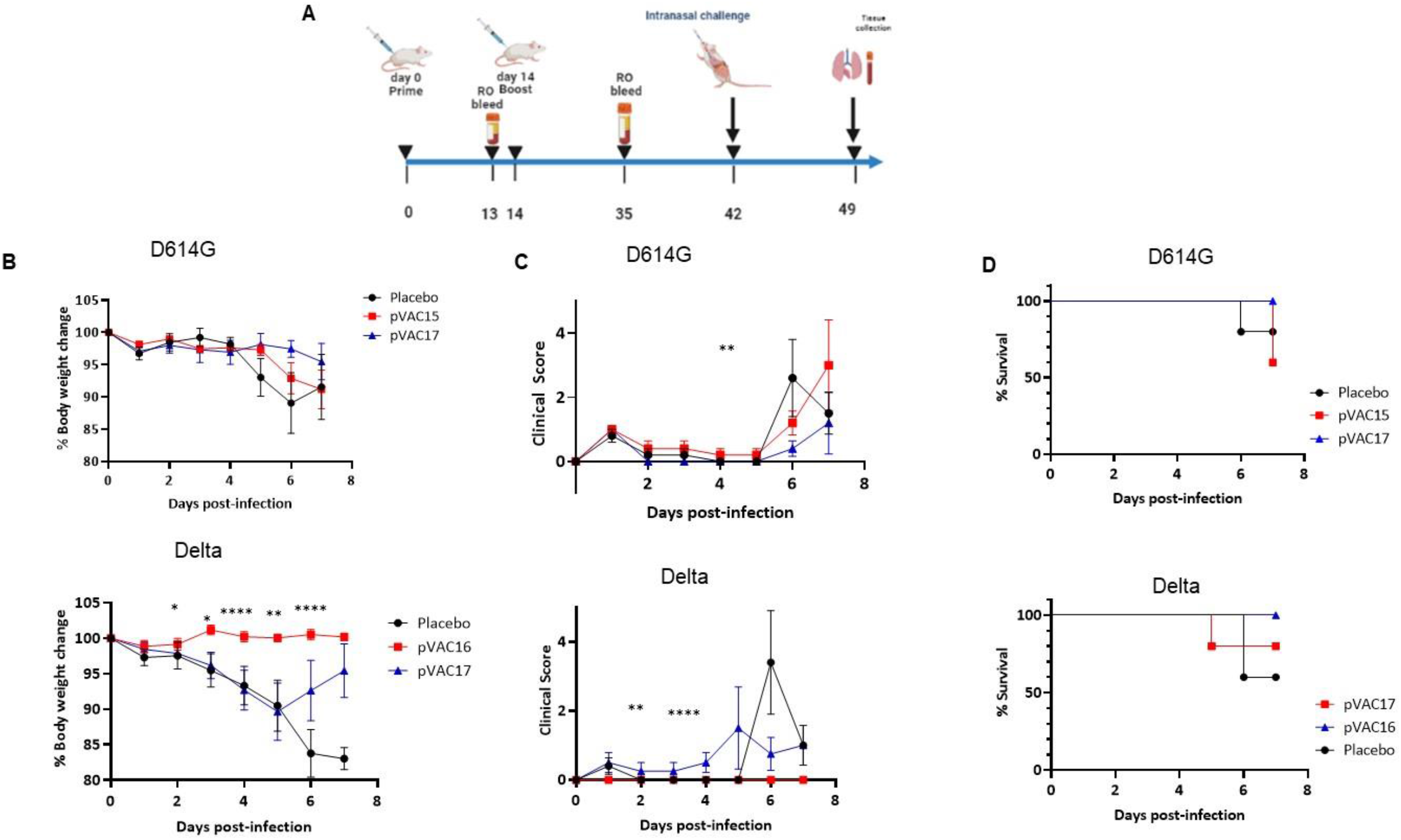
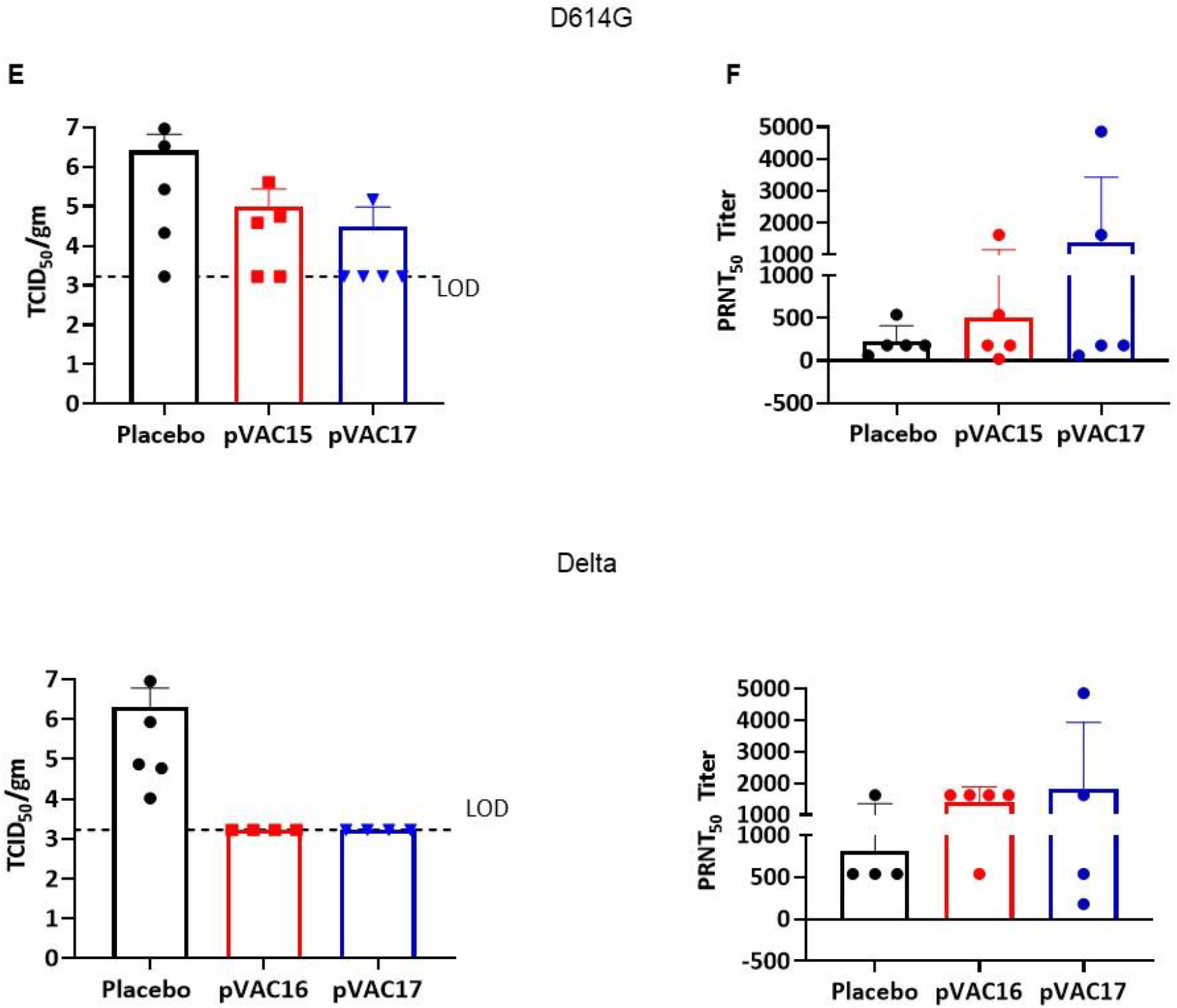
Intramuscular plasmid DNA vaccination protects K18-hACE2 transgenic mice against SARS-CoV-2 infection. (A) K18 hACE2 transgenic female mice were vaccinated with placebo, pVAC15, 16 or 17 with prime and boost, and then challenged intranasally with either SARS-CoV-2 D614G variant or Delta variant (n=5) and monitored for 7 dpi and necropsied. (B) Body weight change. (C) Clinical score based on a standardized 0 to 4 grading system that describes the clinical wellbeing of mice. (D) Mortality of mice. (E) Viral load in lung at 7 dpi. Data shown are in mean ± SD. (F) FRNT IC50 titer of sera from 7 dpi against Delta variant. Limit of detection = 1667 TCID50/gm. P Values were calculated using two-way ANOVA: *P< 0.05, **P < 0.01, **** P<0.0001.

Delta variant virus infection in the Placebo group showed rapid weight loss with a maximum weight loss of 18% at 7 dpi. The pVAC16 immunized mice did not show any weight loss from infection (**Figure 6C**). The pVAC17 vaccinated groups maximum weight loss was 11% at 5 dpi and then started recovering their lost weight at 6 dpi with an average weight loss of 5% at 7 dpi. Disease clinical score peaked at 6 dpi in the Placebo group had a mean score of 3.5 and 40% mortality. The pVAC16 vaccinated group did not show any clinical symptoms and no mortality (**Figure 6C**). The pVAC17 vaccinated group showed mild clinical symptoms with a clinical score of 2 at 6 dpi and 20% mortality. (**Figure 6D**). Mean lung viral load of 2.01×10^6^ TCID_50_ at 7 dpi were detected in Placebo group whereas all 4 surviving mice from pVAC16 and pVAC17 immunized groups had no detectable virus (**Figure 6E**), correlating with body weight loss, clinical score, and mortality observations.

The neutralizing activity of sera from 7-dpi was evaluated using live virus by FRNT assay against Delta variant. The neutralizing titer IC_50_ Geometric mean of the D614G challenged groups were 180, 237 and 433 for placebo, pVAC15 and pVAC17 respectively. The neutralizing titer IC_50_ geometric mean of the delta challenged groups were 540, 1231 and 1044 for placebo, pVAC15 and pVAC17 respectively (**Figure 6F**). pVAC16 and pVAC17 vaccine groups carrying Delta spike antigen showed higher neutralization titers following challenge as compared to pVAC15 and pVAC17 vaccinated and D614G challenged mice.

### Stability and immunogenicity of formulated pDNA

We evaluated the stability and immunogenicity of DNA formulated with the functional polymer and adjuvant. DNA was formulated and stored at either 4C or at room temperature or at -20C. DNA was analyzed by a gel electrophoresis and through immunization and then assessment of induced antibody response by ELISA. Analysis revealed that the supercoiled DNA content significantly declined over time for samples stored at room temperature. Supercoiled DNA content was significantly decreased by ∼ 50% at 1 month, >70% at 3 months and >90% at 6 months (data not shown). In contrast, the supercoiled DNA content remained unchanged for samples stored at 4°C and -20°C even up to 12 months BALB/c mice were immunized on days 0 and 14 by IM administration with either 125 µg pVAC17 DNA on stability (1, 3, 6, 9 and 12 months) or pVAC17 formulated on day of immunization. Three weeks post the second injection sera was collected and tested by ELISA. The level of immune response showed no significant difference between the 4C stability and freshly formulated samples (**Figure 7A**). The 22C stability samples showed a decrease in the induction of immune response by month 3. The immunology data is reflected of the percent supercoiled as determined by gel electrophoresis.

**Figure. 7.**
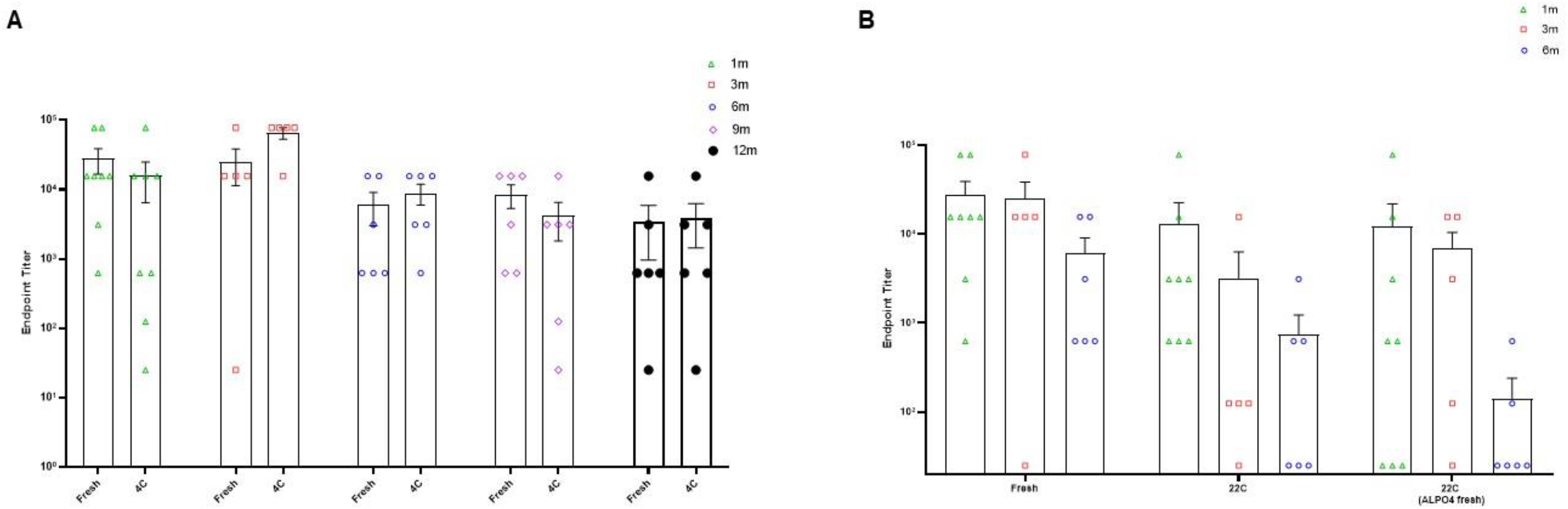
Long term humoral immune response to pVAC17 Sera from immunized with DNA stored at. (A) 4C or (B) 22 C were mice were tested in a standard ELISA. The end point titers to spike full length protein were defined as the reciprocal of the highest dilution that emitted an OD exceeding 4 times the background.

## Discussion

Vaccination against COVID-19 stemmed the pandemic, but also identified areas for improvement. DNA vaccines hold several advantages as both a primary vaccine and as a booster for SARs-CoV-2. For example, the DNA plasmid-based vaccines provide highly durable antigen exposure, elicit strong cellular immune response, have good stability at workable temperatures, offer flexible vaccine design incorporating multiple antigens in a single plasmid to improve vaccine breadth, and have a simple and cost-effective generic manufacturing process. Despite the inherent advantages the progress in DNA-based vaccine has been slow due to inefficient delivery. Viral vectors and devices have been used to improve delivery but are subjected to certain limitations including safety and repeatability with viral vectors and compliance with devices. In this manuscript we describe a DNA-based vaccine approach that is independent of viruses or devices and relies on a functionalized synthetic polymer for DNA plasmid delivery. We demonstrate immunogenicity and protection against SARS-CoV-2 using three different SARS-CoV-2 DNA vaccines formulated with the functionalized polymer and delivered by IM injection in mouse models. All vaccines (pVAC15, pVAC16, and pVAC17) induced humoral and cellular responses that resulted in suppression of viral replication in a mouse model.

The functionalized polymer formulation that was used to test the immunogenicity of pVAC15, pVAC16, and pVAC17 was initially evaluated and optimized for gene transfer using a luciferase plasmid following intramuscular administration. The functionalized polymer improved the luciferase gene expression by 10-fold above the naked DNA (**Figure 2**). In addition to the higher levels of expression of luciferase observed *in vivo,* luciferase expression was observed at background levels for up to 85 days despite co-expression of highly immunogenic spike protein (**Figure 2D**). This long level of expression is analogous to that reported for adenoviral vectors. It has previously been demonstrated that adenovirus-vector DNA can remain detectable for months after injection [39]. Longer duration of antigen expression may be the reason for longer lasting immunity [39] when compared to mRNA [40,41] which is degraded soon after immunization. Pre-existing and vaccination-induced immunity against the vector are a unique feature of adenovirus vector vaccines [42–46]. Therefore, it is unclear if anti-vector immune responses will allow for COVID-19 booster vaccinations with recombinant viral vectors. None-the-less, longer expression can lead to maturation of immune responses particularly antibody somatic hypermutation and thus higher affinity antibody.

DNA vaccines also afford delivery of multiple antigens in a single plasmid DNA molecule without the burden of manufacturing multiple proteins such as in the case of subunit vaccines. Other vaccine platforms have had success delivering multiple antigens such as a VSV-based vaccine [47] and adenoviral vaccine [48]. In the studies presented here pVAC17 demonstrated expression of both SARS-CoV-2 variants D614G and Delta, which resulted in the induction of cross neutralizing antibodies and protection in the mouse model against both a D614G and Delta challenge. Other approaches to targeting multiple VOCs include multiple spikes or RBDs. Zhang was able to induce robust neutralizing immune responses to two variants of concern with a single vaccine [49]. DNA vaccines could in fact not only encode antigens to target two viral strains, but with the incorporation of molecular sequences such as furin cleavage sites a series of neutralizing epitopes can be incorporated into a single vaccine backbone. Such is the case in the area of cancer vaccines where greater than 10 neoantigens can be encoded into a single plasmid [50].

In summary, these results demonstrate that DNA can be delivered with a functionalized polymer that leads to long term *in vivo* expression and induction of robust protective immune responses. The PlaCCine formulation allows for device free delivery requiring only an IM needle injection. The immunogenicity of PlaCCIne vaccines described in this manuscript have been demonstrated in non-human primates (manuscript in preparation). And a human clinical trial of PLACCINE vaccine is an important next step.

